# Mating behaviour and connectivity drive sporadic European bat lyssavirus 2 infections in *Myotis myotis*

**DOI:** 10.1101/2025.09.22.677820

**Authors:** Frédéric Touzalin, Marine Wasniewski, Evelyne Picard-Meyer, Olivier Farcy, Yann Le Bris, Arnaud Le Houedec, Corentin Le Floch, Megan E Griffiths, Sébastien J. Puechmaille, Emma C. Teeling, Juan Morales, Mafalda Viana, Daniel G. Streicker

## Abstract

Understanding the transmission and persistence of zoonoses in wild reservoirs is crucial for outbreak prevention, demographic assessment, and conservation planning. Host demography, ecology, behaviour and within-host viral kinetics together underpin transmission dynamics but are rarely united within epidemiological models. Here we characterised the transmission dynamics of European bat lyssavirus 2 (EBLV-2) infection dynamics by uniting a long-term study (2010-2018) of the ecology of five greater mouse-eared bat (*Myotis myotis*) colonies in France (∼2700 marked individuals) with diagnostic evidence of virus circulation. Although no lyssavirus was detected in any of the 442 saliva samples tested, seropositivity showed synchronised peaks across colonies (0-100% annually), with the highest seroprevalence in 2015-2017, but remaining low otherwise. Fitting Bayesian compartmental epidemiological models, we found evidence that shifts in EBLV-2 prevalence were primarily driven by the gregarious behaviour of populations during the mating season and age-dependent variation in transmission. The duration of maternal immunity emerged as a key, and still empirically unknown factor, which shaped whether EBLV-2 maintenance was enzootic or required an external force of infection. *Lyssavirus* outbreaks in *M. myotis* were associated with few deaths overall and had a negligible impact on population dynamics. These findings underscore the value of integrating ecological and individual-level data with mechanistic modelling to enhance our understanding and management of zoonotic pathogens in wildlife.

## Introduction

Transmission of pathogens from wildlife to humans (zoonotic transmission) is a major source of newly emerging infectious diseases (EIDs) [1]. Predicting and preventing cross-species pathogen spillover is a global priority but remains challenging [2]. The mechanisms that sustain most zoonotic EIDs in reservoir hosts remain poorly understood, complicating surveillance and prevention [3]. Longitudinal serological surveys coupled with data-driven mechanistic modelling offer a promising way forward to explicitly test biological and ecological hypotheses of disease maintenance and transmission and to infer temporal patterns of spillover risks [4]. However, the lack of ecological, biological, and behavioural data on reservoir hosts remains a widespread impediment to realistic modelling of the transmission dynamics of zoonoses in natural populations. *European bat lyssavirus 2* (EBLV-2) is a member of the zoonotic *Lyssavirus* genus and one of 15 bat-associated lyssaviruses among the 18 currently recognised species within the family *Rhabdoviridae* [17]. EBLV-2 is the second most detected *Lyssavirus* circulating in Europe, following *European bat lyssavirus 1* (EBLV-1) [18,19]. Before this study, it was only identified in Daubenton’s bats (*Myotis daubentonii*) [8–10] and in pond bats (*M. dasycneme*) [6,11], mainly distributed in the north of Europe, including the United Kingdom, the Netherlands, Germany, Switzerland, Finland, Norway and Denmark [8,9,11–15]. Since its identification in 1985, tens of cases have been confirmed [16], mainly involving bats, except for two fatal human cases, due to bat bites, in Finland in 1985 and in Scotland in 2002 [33]. Natural spillover to other non-bat species has not been reported, and experimental infections indicate low susceptibility in sheep and foxes [19,20]. Most cases have been detected through passive surveillance, predominately involving dead or morbid bats found in human-inhabited areas, leading to species representation bias and regional variation in case reports. Active lyssaviruses surveillance in wild European bats remains rare and is spatio-temporally restricted to a few maternity colonies, e.g., EBLV-1 in Spain and France and EBLV-2 in the United Kingdom [10,21,22].

Key aspects for EBLV-2 transmission in bats remain poorly defined. An experimental study in Daubenton’s bats suggested the EBLV-2 infection can cause weight loss, paralysis and death indistinguishable from rabies, but most exposed bats failed to develop clinical disease or shed virus, most likely representing ‘abortive’ infections [23]. A field study in free-ranging bats of the same species suggested that mixing of bats between roosts during seasonal swarming events might enable EBLV-2 maintenance [10]. However, it is increasingly evident that persistence dynamics of *lyssaviruses* depends both on the specific virus involved and the ecology of the putative reservoir host species. For example, vampire bat (*Desmodus rotundus*) rabies appears to be maintained by spatial processes such as metapopulations or invasion fronts [24], insectivorous bat (*Eptesicus fuscus*) rabies in temperate North America persists through seasonal pulses of transmission after hibernation [25] and European bat lyssavirus-1 (EBLV-1) relied on immune waning during hibernation and seasonal birth pulses to persist in *M. myotis* bats in Italy [4]. These studies highlight how the impacts of immunity, seasonality, social behaviour, or movements between colonies can influence viral persistence but require assumptions which could largely not be verified independently with empirical data. For example, the absence of longitudinal sampling of individually marked bats precluded using seroconversion histories to inform exposure rates or waning immunity, an approach that has proven powerful in other bat-virus systems [26]. Similarly, seasonal movement patterns and connectivity between bat roosts have been assumed, rather than informed by observational or animal tracking data. Finally, bat population sizes and demographic rates are rarely known with precision, obscuring the ecological realism of models [24,26]. To date, no study on *Lyssavirus* epidemiology has successfully incorporated both detailed understanding of host demography/behaviour and within-host viral kinetics from longitudinally sampled individual bats.

Here, we used ecological and molecular longitudinal data collected from 2010 to 2018 across five greater mouse-eared bat (*Myotis myotis*, Vespertilionidae) maternity colonies in France to develop data-driven Bayesian integrated epidemiological models for EBLV-2. We first employed mixed-effects logistic regression to identify drivers of lyssavirus seroprevalence. Then, we used Passive Integrated Transponder (PIT) monitoring to characterise the seasonal patterns of colony distribution and connectivity and identify risk factors for inter-colony transmission and external introductions. Finally, by integrating demographic and molecular data in a mechanistic model, we estimated key epidemiological and demographic parameters, offering new insight into EBLV-2 dynamics and informing more targeted public health strategies.

## Materials and Methods

### Serological data

Blood samples were collected between 2010 and 2018 in five *Myotis myotis* maternity colonies in south Brittany: La Roche Bernard (47◦ 31’N, 2◦ 18’W), Férel (47◦ 28’N, 2◦ 20’W), Noyal-Muzillac (47◦ 35’N, 2◦ 27’W), Béganne (47◦ 35’N, 2◦ 14’W), and Limerzel (47◦ 38’N, 2◦ 21’W). These colonies are < 20km apart and situated at the north-western limit of the species’ range in Europe. Bats emerging from the colonies were sampled once a year, between the end of June and early July, after juveniles started to fly out of the roost. Captures were carried out in accordance with the French ethical and sampling guidelines issued in 3 successive permits delivered by the Préfet du Morbihan (Brittany, France) to O. Farcy and F. Touzalin. The capture was carried out using custom harp-traps placed in front of the exits. After verifying the presence of PIT tags, specifically ID-100C (Trovan®) with a unique 10-digit code (2.12×11 mm, 0.1gr), bats were placed in individual cloth bags. Bats not previously tagged received PIT tags and all individuals underwent biometric assessment. From a subset of bats, blood was taken at the inter-femoral vein, located in the uropatagium along the posterior limbs, by piercing the vein with a sterile 27G needle. The blood drops were adsorbed onto a maximum four Whatman paper discs (cut from Mini Trans-Blot® filter paper, Bio-Rad) to contain each 20 µL of whole blood once fully impregnated. Bleeding was stopped using Stop Hemo© powder or Hemostop© gel. After soaking, the Whatman paper discs were dried at room temperature and then stored at –20°C until serological analysis. Antibody titration for EBLV-2 and EBLV-1 was conducted in parallel by the French Agency for Food, Environmental and Occupational Health & Safety (ANSES), Laboratory for Rabies and Wildlife, located in Nancy (France); see electronic supplementary material 1 (section 1.1). We considered detectable antibody titres to indicate protection, which corresponds to individuals who seroconverted and survived after exposure to the EBLV virus, although there may be cases where protection from virus neutralising antibodies is not guaranteed or, conversely, where protection is effective despite the absence of detectable titres following multiple exposures [27].

### Molecular detection of lyssaviruses in bat saliva

To assess if animals were shedding lyssavirus, 442 saliva samples were collected using commercial sterile flocked nasopharyngeal swabs (Ø 2.5 mm head) and placed in RNA-later solution for storage. Reverse transcription polymerase chain reaction for the detection of lyssavirus RNA was carried out by ANSES (see electronic supplementary material 1, section 1.3).

### Risk factors for EBLV-2 seropositivity

We fitted a Bayesian logistic regression model, including a joint framework to estimate missing bat colony counts, to investigate predictors of EBLV-2 seropositivity. The model incorporated demographic factors (sex, age, and a sex × age interaction to account for unbalanced sampling), colony size (total number of adults and juveniles), biometrical variables (forearm length, weight, and the Scaled Mass Index, SMI, as a measure of body condition [28])), sample size, and the reproductive status of adults. Based on the correlation matrix of the covariates (figure S3, electronic supplementary material 1), we excluded adult female and juvenile counts because both were highly correlated with total colony size (r > 0.8). Reproductive status comprised males, all sexually inactive at the time of sampling, and females classified as lactating, non-lactating (non-reproductive that year), or post-lactating (table S1, electronic supplementary material 1). Colony was included as a random effect, assuming more homogeneous exposure and transmission risk within colonies than between them. To assess spatio-temporal synchrony in seroprevalence among maternity colonies across years, we further included year and the year × colony interaction as random effects and calculated the intra-class correlation coefficient (ICC). The ICC was calculated as the ratio of the variance attributable to the temporal random effect (year) to the sum of the variances of the temporal (year) and spatio-temporal (year × colony) random effects. This metric quantifies the degree of synchrony among colonies through time, with values approaching one indicating high spatio-temporal synchrony (electronic supplementary material 1, section 1.5). Finally, we compared four models that differed in the probability distribution used to estimate missing colony counts. Specifically, we evaluated a Poisson distribution and three negative binomial parametrisation, in which the dispersion parameter was either shared across all observations, colony-specific, or allowed to vary by both colony and age class within each colony (figure S5). To run the models, we used the package jagsUI [29], an interface for jags [30] in R 4.4.3 [31]. Four chains, with 500,000 iterations each, including a burn-in phase of 200,000 iterations. Chains were sampled every 600 iterations and model convergence was checked using the Rhat statistic and effective sample size.

### Seasonal field data collection

We analysed data collected throughout three phases of the annual biological cycle, (1) gestation and rearing, (2) mating, and (3) wintering (figure 1.b), to investigate bat demography and connectivity patterns among maternity roosts and assess their potential epidemiological implications. Yearly counts and capture events were organised with Bretagne Vivante-SEPNB, a regional conservation NGO, that has monitored maternity roosts and wintering sites in Brittany for more than two decades. In early June, prior to juvenile fledging, adult bats were counted as they exited the roost through direct observation. After adults’ emergence, juveniles were counted inside the roost, either by direct observation for small groups (classically <50) or by analysing photographs for larger groups. During the gestation and rearing (∼16 Feb. to 31 Aug., figure 1.b), we captured bats at maternity colonies using mist nets (end of June/early July) for sampling and marking. The colonies were equipped with antennas and stationary PIT tag recorders, allowing individuals to be detected all year round when entering/exiting roosts. For technical reasons, no PIT tag reader was deployed at the Limerzel colony. Additional captures at foraging sites (woodlands) enabled assessment of home-range overlap among colonies. During the mating season (∼1 Sept. to 15 Oct.) bats were captured at underground sites, often called swarming sites, and checked for PIT tags. After exploring tens of potential mating sites, we limited our annual monitoring of mating site use to two sites that showed a substantially higher capture rate of *M. myotis* bats than all the others. During hibernation (∼16 Oct. to 15 Feb.), ∼30 hibernacula were checked three times per winter without handling bats to minimise disturbance, using hand-held antennas.

**Figure 1.**
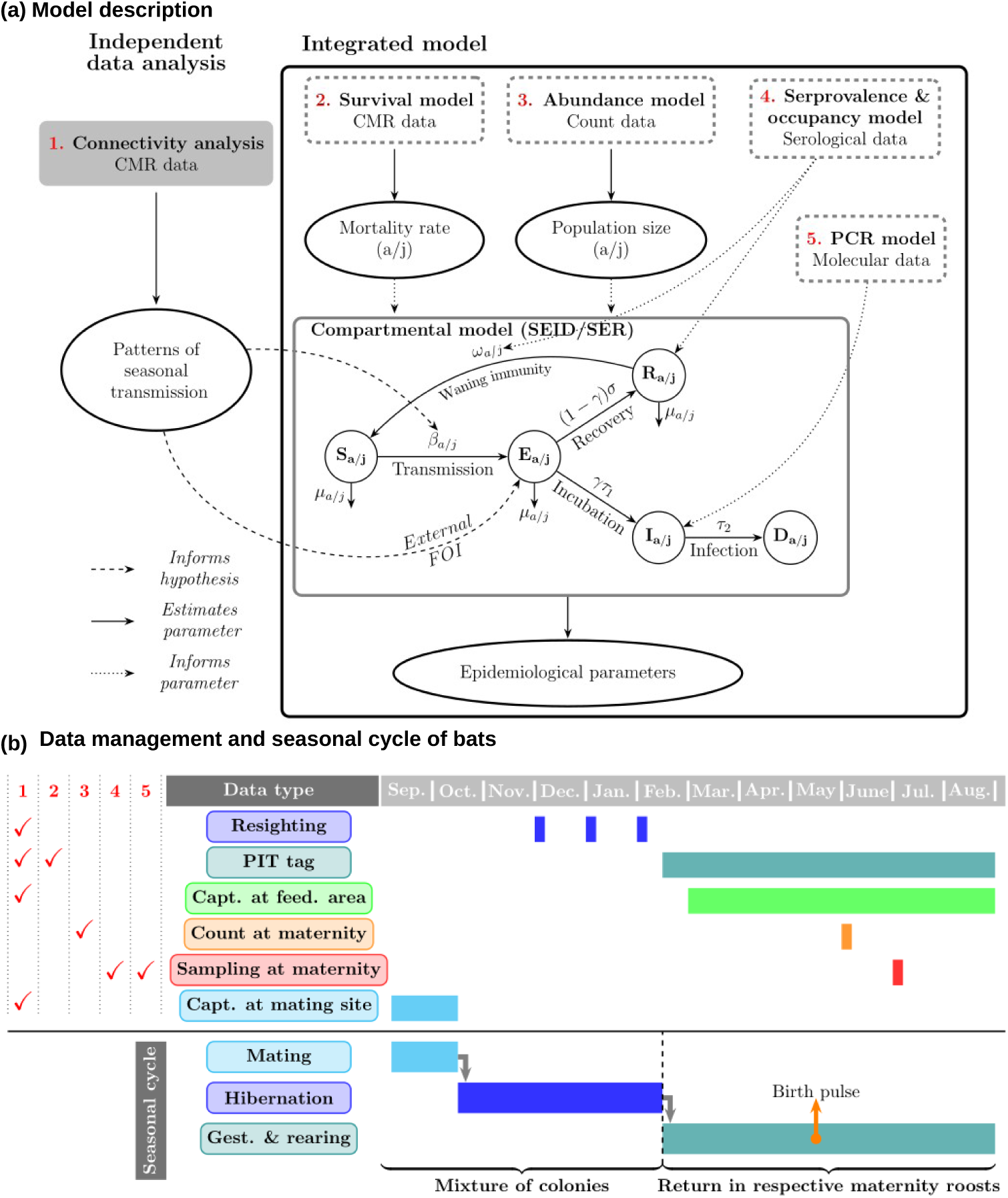
Biological and mathematical description of the study system. **(a)** Schematic description of the model. Connectivity analyses (1) provide insights into behavioural ecology and help narrow down the hypotheses regarding epidemiological mechanisms to be tested, particularly those related to seasonal transmission risk and periods for potential importation of EBLV-2 infection from unstudied populations. Integrated Bayesian model: survival model (2) estimates mortality rate, and abundance model (3) estimates population fluctuation through time, both models informing the compartmental model. The seroprevalence data and the occupancy model on repetitively sampled individuals (4) depend on the proportion of recovered individuals (R), and the binomial logistic regression (5) on PCR data (detection of the virus) depends on the proportion of infected individuals (I). Compartmental model: susceptible individuals (S) become exposed at a transmission rate β, γ is the fraction of exposed bats (E) that become infectious (I) at a rate τ_1_ and 1 – γ the fraction that seroconvert and recover (R) at a rate σ. All infected individuals die at a rate τ_2_. μ is the rate of mortality due to causes other than lyssavirus infection. Depending on the scenario (see Hypotheses tested in the main text), an external force of infection is allowed from immigrated infectious individuals, which can increase the number of exposed individuals. the subscript “a/j” indicates parameters estimated separately for adults and juveniles. **(b)** Data collection period by data type and their use in the analyses (in red), as indicated by the notation in figure part (a). Passive tag search occurred during winter (3 sessions/year) and at maternity colonies (stationary recorders), capture sessions occurred at foraging grounds, maternity (for blood sampling – early July) and at mating site. Bat counts were carried out at maternity colonies. The annual life cycle of M. myotis bats is defined as: gestation and rearing season (mid-February, to August), with females settling at the maternity colonies (buildings), birth pulse occurring mid-May, mating (September to mid-October) and hibernation seasons (mid-October to mid-February), when all colonies of bats leave the maternity roosts and gather at various underground sites (usually mines).

### Spatio-temporal connectivity

Geographic locations of capture at foraging grounds were analysed using adaptative local convex hull and kernel density approaches to estimate home-range overlap among maternity colonies. Capture-mark-recature (CMR) data, based on PIT tag records, from both captures and stationary readers at maternity roosts were used to assess individual movements between colonies within gestation and rearing for adults and juveniles. During the mating season, colony connectivity was evaluated by counting instances where marked individuals from different maternity colonies were detected on the same night. A similar analysis quantified cases where individuals from multiple maternity colonies were observed in the same hibernacula. Given the sufficient number of hibernacula, adaptive local convex hull and kernel density approaches were also applied to estimate winter home-range overlap. All analyses were conducted in R 4.4.3 [31], employing the adehabitatHR, MASS, sf and igraph packages. The packages ggspatial, tidygraph and ggraph were used for graphical representation [32–39]. All codes can be found in the electronic supplementary material 1.

### Demographic parameter estimation

Overall, CMR data used in this study were a combination of capture data and records from stationary/hand readers. However, since 18% of juveniles and 10% of adults may have lost PIT tags [36], we also used genotype, based on 16 microsatellite markers (see description in [40]), as a genetic marker to recover individual identities despite tag loss. Mortality rates were assessed using a two-age-class Cormack-Jolly-Seber (CJS) model [41–43] in a multinomial likelihood framework [44,45]. To address the identifiability issue in the final survival interval, where survival and detection probabilities are confounded, the analysis was extended to data collected between 2010 and 2019. These annual estimates encompassed all mortality causes for both age classes, including EBLV-2 infection. Population dynamics were inferred from count data. To account for uncertainty and missing values in colony counts, we applied the same Poisson regression model used in the logistic analysis described above, incorporating a cubic polynomial time effect to capture inter-annual variations in the effective numbers of adults and juveniles (electronic supplementary material 1, section 1.5).

### EBLV2 transmission dynamics

We used Bayesian inference to fit mechanistic compartmental models to the longitudinal EBLV-2 seroprevalence data (figure 1.a). Since the risk factor analyses detailed above indicated synchrony in seroprevalence among these colonies (see results and figure 2), and given the limited data available for single colonies, we pooled the data across the five colonies, treating them as a single population. Our base model followed a SEID/SER structure, which assumed two possible fates for the bats exposed to EBLV-2: recovery from exposure without ever becoming infectious or death following a period of infectiousness, as commonly assumed in observational field studies and supported by *in vivo* challenge studies [4,23–25,46,47]. In this model, a fraction (γ) of exposed bats (E) become infectious (I) at a rate *τ1* and then die (D) at a rate *τ2* while the remaining exposed bats (1-*γ*) recover at a rate *σ* with immunity (R) for a period 1/*ω* without being infectious (figure 1.a, electronic supplementary material 2, section 1). Compartment D serves to estimate the number of animals succumbing specifically to infection-related mortality, distinguishing these cases from deaths attributable to other causes. We modelled the population as a two-class, age-structured system, where juveniles were defined as newborns up until the following spring, at which point they had matured into adults and a fraction began reproduction, as supported by data on reproductive statuses during capture. We also accounted for the biological seasonality of the studied species in Brittany (figure 1.b), based on the observed data and CMR records. In particular, a synchronised birth pulse was assumed to occur in mid-May, which was deduced from observations during counts that occurred annually in early June. Each year, a fraction of new-born individuals (juvenile cohort)were considered protected against infection by maternal immunity (M) in the same proportion as the seropositive females (R), while the other fraction were assumed susceptible (S). The duration of maternal immunity is crucial for infection mitigation, as seasonal birth pulses have been considered a key factor in the persistence of lyssavirus infections [4]. We then defined two possible durations (short/long) for this parameter, described below. Mating season, during which bats leave the maternity colonies and congregate at mating sites on a large geographic scale, was considered the timing when infection can be imported, i.e., by an external force of infection (FOI). This implied contact with infected bats from other maternity colonies outside the studied system or other bat species (figure 1.a).

**Figure 2.**
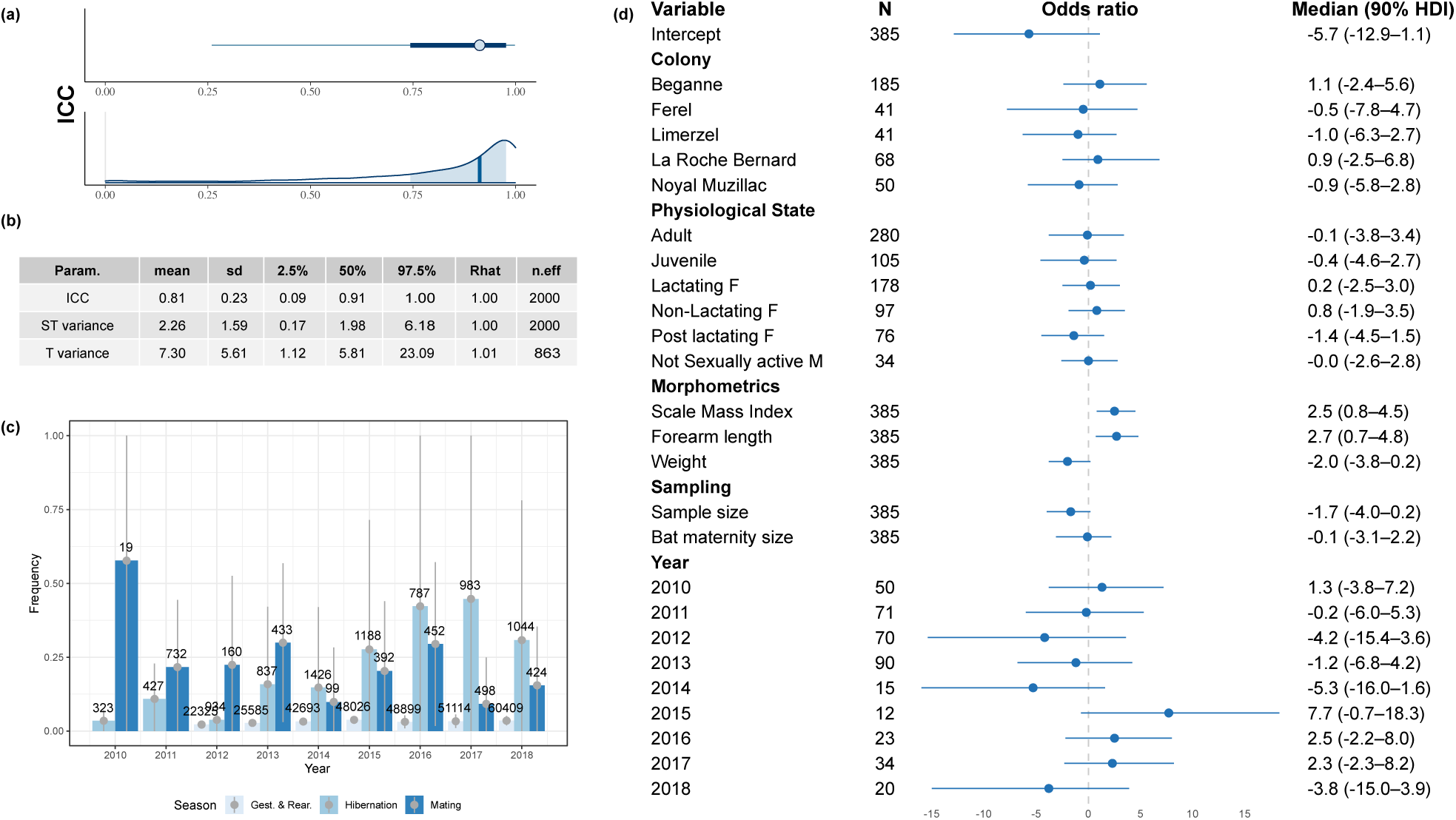
Drivers of EBLV-2 serological dynamics in M. myotis in France. **(a)** Top, median (dot), 90% probability mass (thin line) and 50% probability mass (thick line) of the intra-class correlation coefficient (ICC). Bottom, posterior distribution with median (blue line) and 50% probability mass (light blue area). **(b)** Logistic regression, posterior summary for ICC, spatio-temporal variance (ST), and inter-annual variance (T). **(c)** Annual frequency of individuals (bat/day) from different maternity roots observed in the same site, calculated at three different seasons. Numbers at the top of the bars indicate the cumulative number of bat/days recorded across each season. During gestation and rearing season, RFID records were used, while capture data were used during mating season and resightings during winter. **(d)** Median (dot), 90% probability mass (thin line) of the logistic regression parameters. For each parameter, the number of observations (N), the mean value and the 90% highest (posterior) density interval (HDI) are indicated.

Compartments S, E, and R were considered susceptible to causes of death unrelated to lyssavirus infection at a rate *μ*. We assumed that infectious individuals (I) succumb only from the infection itself, as both experimental data and field observations indicate that the period of infection preceding death is brief (1-2 weeks). The death rate *μ* was estimated as the total mortality rate (estimated by the CJS model) minus the death rate attributable specifically to lyssavirus infection (compartment D).

For the first year of the study, we used priors to estimate the initial proportions of the E, R, and I compartments. These priors were defined with density distributions concentrated near zero and hierarchically defined to ensure that their sum did not sum to one. We used informative priors when published studies or expert knowledge from bat rescue centres were available (table S1; electronic supplementary material 2). The differential equations of the compartmental model are detailed in the supplementary material 2 (section 1.2).

Integrating multiple data streams in epidemiology, such as combining ecological, individual-level, and serological data, enhances model accuracy and insight, contrasting with most traditional models that rely solely on a single data source like population-level serology [26,48]. Five likelihoods were integrated into our model, all collectively informing the SEID/SER framework (figure 1.a): (1) CMR data; (2) abundance of adults and juveniles; (3) population-level seroprevalence; (4) individual seroconversion; (5) PCR data. The survival (CJS) model estimated the total mortality rate, while the abundance model provided annual estimates of adult and juvenile population sizes. Serological data were used initially to inform recovery rate by modelling seroprevalence as a binomial outcome with the probability corresponding to the proportion of recovered individuals (R). Subsequently, the serological data collected from individuals recaptured multiple times were integrated into an occupancy model to inform the immune period (ω) by predicting the probability of seroconversion or reversion between two sampling points [26]. Finally, molecular (PCR) data were employed to inform the infection rate by estimating the presence or absence of viral particles as a binomial outcome, where the probability reflects the proportion of infected individuals (I). To quantify the relative contribution of each age class to transmission over the study period, we computed the time-averaged age-specific effective reproduction number:

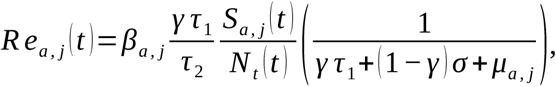

with subscripts “a,j” corresponding to adults or juveniles, respectively (see section 1.3 in electronic supplementary material 2).

### Alternative epidemiological models

Despite our intensive exploration of bat behaviour from our CMR data, certain biological phenomena regarding transmission dynamics remained unresolved. Consequently, we defined various scenarios to test the biologically plausible alternative hypotheses. In scenario 1 (maternal immunity duration), we proposed either that antibody detection in juveniles at the time of sampling (around 8 weeks of life) would indicate (1) recovered from lyssavirus exposures early in life if maternal immunity is short (four weeks’ scenario) or (2) the presence of maternal antibodies if maternal immunity persists for more than 8 weeks (twelve weeks’ scenario). In a second scenario (transmission during winter), we assumed that in winter, due to torpor, bats (1) had no interactions, with all rates (*β, γ, τ1, τ2, σ, ω*) and the mortality rates set to zero, or (2) as CMR data indicated bat movements during winter, all rates were halved during winter hibernation, except for transmission rate. Transmission was estimated from the data with a uniform prior with a range of 0 to half its value during the active period to account for the rare interactions during this period. For the third scenario (transmission dynamics before winter), we proposed that either (1) *β* remained constant outside wintering period or (2) *β* increased with a sigmoid shape, starting at a minimum value from hibernation arousal and reaching a maximum by the end of the mating season. This scenario was inspired by the increase in interaction along the breeding season with the birth pulse and mixing of population during the mating period. In a fourth scenario (transmission rate dependence), we considered either (1) frequency or (2) density dependence in transmission dynamics. Scenario five (external FOI) proposed that importation events may occur (1) sporadically, driven by transmission processes operating at broader geographic scales beyond the study area or by other bat species, or (2) cyclically, which may account for alternating periods of low and high seroprevalence as observed in foxes [49], or (3) that EBLV-2 is enzootic (i.e., no importation is required from external sources). The external FOI (Imp) was modelled as a per-susceptible hazard function, a proportion of susceptible individuals that can be exposed by external sources (electronic supplementary material 2). A final scenario (age-class transmission rate) tested heterogeneity in transmission between age classes, with either (1) a similar or (2) a different transmission rate between juveniles and adults. Integrated models were run in jags using the jagsUI interface for R [35]. We used four MCMC chains, with 500,000 iterations each, including a burn-in phase of 200,000 iterations. One sample out of 600 was saved (2000 samples per parameter) and model convergence was checked using the Rhat statistic and effective sample size. A set of 96 models was run to test all possible combinations of scenarios, which were then ranked using the Leave-One-Out Information Criterion (LOOIC), difference in expected logarithmic predictive density (ELPD) and two model-weight approaches, stacking and Pseudo-BMA+ [50].

## Results

### Population and individual level patterns of seropositivity

A total of 442 blood samples were collected from five maternity roosts of *M. myotis* bats in Brittany among 23 sampling events between 2010 and 2018. Only 399 had enough volume (60-80uL of whole blood) to allow the mFAVNt test to be performed for both EBLV-1 and EBL-2 titres. Across all roosts and time points, 10% (n=40) of these samples tested positive for antibodies against EBLV-2, none for EBLV-1. Seroprevalence exhibited pronounced fluctuations over time, ranging from 0 to 100% (figure 3; electronic supplementary material 1, figure S1), with seroprevalence close to zero during the first 5 years and sudden peaks from 2015 to 2017 followed by a decrease in 2018. According to the LOOIC, the Bayesian logistic regression model with a negative binomial distribution and colony-specific dispersion parameter received the strongest support. However, differences among models support were small, indicating that no single model was clearly superior (table S2, electronic supplementary material 1). We therefore retained the Poison model, given the smaller number of parameters, for subsequent analyses. The ICC was estimated to be close to one, indicating a high degree of spatio-temporal synchrony among roosts. There was strong evidence for a positive effect for forearm length (mean 2.6, 95% HPDI [0.7, 4.8]), the SMI index (mean 2.5, 95% HPDI [0.8, 4.6]), likely reflecting that most seropositive individuals were adult females, who are larger than males and juveniles. Seropositivity was not significantly influenced by age, sex, colony, sample size, or reproductive status (figure 2, electronic supplementary material 1, figure S6). Strong and geographically synchronised inter-annual variation in seroprevalence suggests viral circulation between the maternity colonies in our study area.

**Figure 3:**
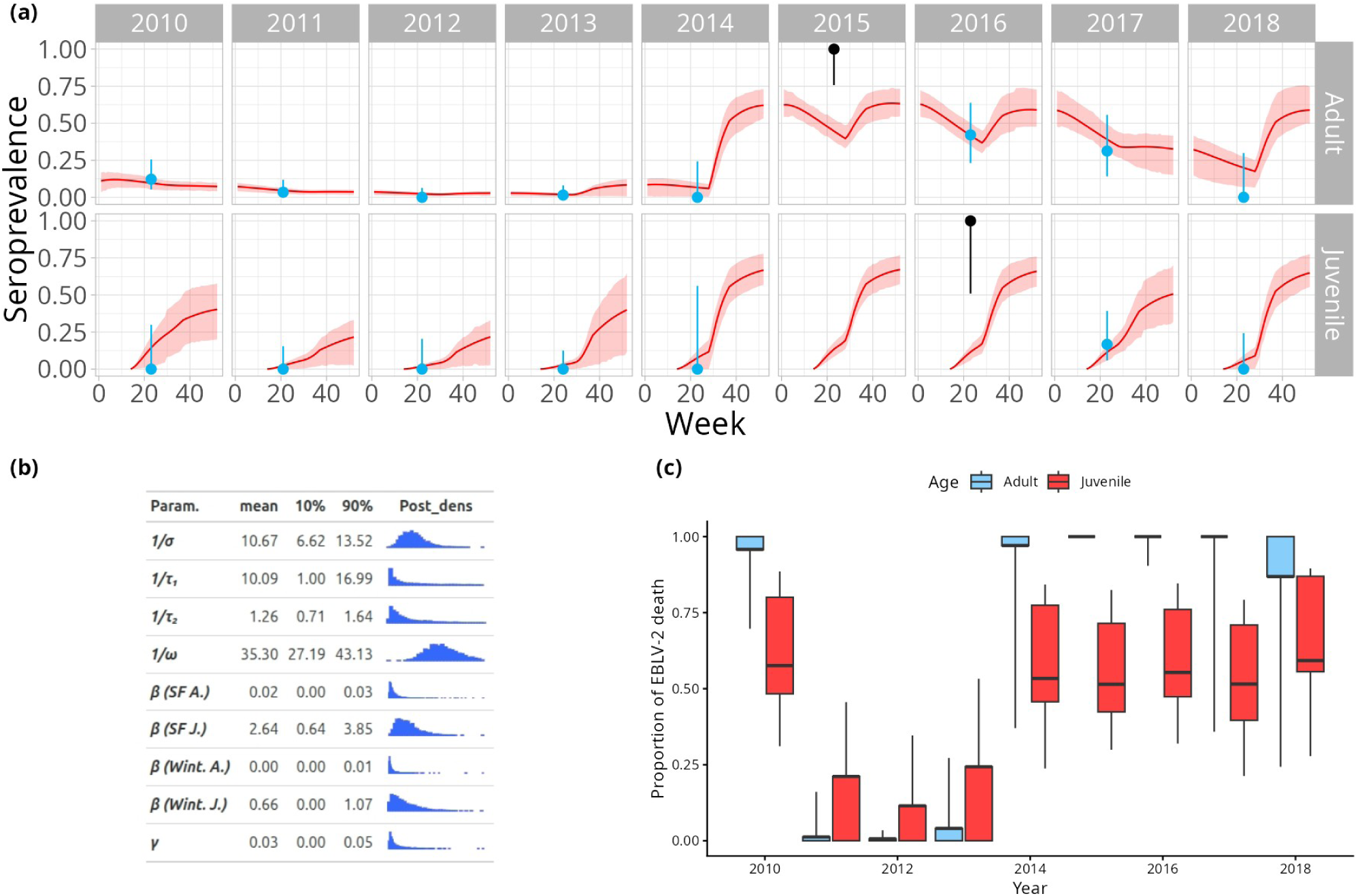
Posterior predictive distribution of the most strongly supported model among the nine candidate models. **(a)** The red lines show the mean predicted trajectory for the recovered class from the best model. Shaded regions show the 80% HPDI. Blue dots show observed population-level seroprevalence data with binomial confidence intervals. Observations that fell outside the 80% HPDI of the posterior cumulative distribution are shown in black. **(b)** Estimated parameters of the compartment model showing posterior means, 80% HPDI and posterior distribution. Transmission rates (β) are given for adults (A.) and juveniles (J.) during spring-to-fall (SF) and winter (Wint.), along with the proportion of individuals who develop infection after exposure (γ). The recovery period (1/σ), incubation period (1/τ_1_), infectious period (1/τ_2_) and immune protection period (1/ω) are reported in weeks. **(c)** Proportion of death specifically due to EBLV-2 infection in both age classes. The thick horizontal line marks the mean, the sides of the box denote the 50% quantile range, and the whiskers extend to the 10th and 90th percentiles, encompassing the central 80% of the distribution.

A total of 45 individually marked bats were recaptured and sampled at least twice during the study period (range: 2-3 captures, time between sampling: 1-7 years). Of 51 sample pairs, 7.8% were seropositive at both time points, 13.7% seroconverted (0-1) and 5.9% showed complete antibody waning (seroreversion). Among bats captured >2 times, three cases of seroreversion (1-0) were observed (electronic supplementary material 1, figure S2), in one case in a one-year interval. These results provided direct evidence of antibody waning over time, as documented in related *Lyssaviruses* species [46]. All the saliva swabs analysed by RT-qPCR tested negative for any *lyssavirus*.

### Seasonal connectivity between colonies

Despite some inter-colony contacts being observed across all seasons (electronic supplementary material 1, figures S7-S19), mating and hibernation periods exhibited the highest frequency of interactions (figure 2.c and section 8 in electronic supplementary material 1). Indeed, the frequency of mixing between colonies was consistently higher in hibernation and in the mating season across the studied period, despite the lower number of observations compared to gestation and rearing. The mating season, in particular, emerged as a critical window for infection spread, driven by increased contact rates associated with sexual activity and male territorial behaviour [51]. In contrast, during hibernation, torpor substantially reduced opportunities for direct interactions among individuals. These patterns suggest that the mating season represents a key period for EBLV-2 transmission, facilitated both by infected individuals migrating from a long distance for copulation and by species mixing. Overall, transmission risk among colonies appeared to remain low during gestation and rearing phases but increased markedly during periods of extensive population mixing, notably during mating.

### EBLV-2 dynamics in *M. myotis*

Across the 96 candidate compartmental epidemiological models evaluated, only 23 achieved full parameter convergence (electronic supplementary material 2, section 2). Non-convergence in the remaining models likely reflects incompatibilities between model assumptions and the empirical data. Models that did not converge were more likely to assume homogeneous transmission rates across age classes, transmission during winter, and the absence of an external force of infection (i.e., purely endogenous infection dynamics). Notably, among the eight models incorporating all three of these assumptions, only one (12.5%) converged.

Because all models exhibited some observations with Pareto k > 0.7, indicating uncertainty in LOOIC reliability, model comparison relied on a combination of approaches. Specifically, we report both pseudo-BMA+ and stacking weights, which are less sensitive to problematic Pareto diagnostics. Collectively, these metrics consistently identified a single model as outperforming the others. The highest-ranked model incorporated several key features: age-dependent transmission heterogeneity, with higher transmission rates in juveniles than in adults; constant transmission during the active season (spring to fall); transmission during winter; frequency-dependent transmission dynamics; and sporadic case importation during the mating season. Posterior predictive checks indicated good overall model fit, with no major discrepancies between observed and predicted values (electronic supplementary material 2, figure S1–S3). Nevertheless, stacking weights across models were relatively diffuse, suggesting that a subset of models provided comparable predictive performance rather than a single clearly dominant model. This pattern is unsurprising given the limited size of the serological dataset and the subtle structural differences among competing models.

When restricting attention to models contributing at least 5% of the predictive signal (stacking), nine models together accounted for 88% of total model weight. Within this subset, several consistent patterns emerged. Most models (78%) supported age-related transmission heterogeneity and similarly (78%) excluded winter transmission, while only two (22%) were compatible with enzootic infection dynamics, instead requiring an external force of infection. Greater variability was observed in the assumptions regarding transmission outside the winter season, with 67% of the models favouring constant transmission rather than transmission that increases during that season, and an almost equal proportion of models (5 out of 9) favouring frequency-dependent transmission processes versus density-dependent ones. The duration of maternally derived immunity emerged as a critical determinant of model behaviour. When maternal immunity was long (four months), the data were best explained by sporadic reinfection driven by the external FOI during mating season. In contrast, shorter maternal immunity (around four weeks) allowed either enzootic persistence or cyclic reinfection.

Parameter estimates (figure 3b, electronic supplementary material 2, section 5) from the top-ranked model indicated that waning immunity (1/*ω*) persisted for more than eight months (mean 35 weeks; 80% HPDI: 28–45; figure 3b). The period between exposure and recovery (1/*σ*) was estimated at approximately 11 weeks (80% HPDI: 8–17). The incubation period (1/*τ_1_*) averaged around ten weeks but showed substantial uncertainty, which may in part reflect high variation in the incubation period for lyssaviruses in bats (80% HPDI: 2–52). The infectious period (1/*τ_2_*) was short, at roughly one week (80% HPDI: 1–2). Importantly, the proportion of infected individuals progressing to lethal infection (*γ*) was very low, estimated at 3.4% (80% HPDI: 0–5.3). Using the next generation matrix, the mean estimated *R_0_* of EBLV-2 before birth pulse was 6.10^-4^ (80% HPDI: 2.10^-4^–9.10^-4^), after birth pulse was 0.1 (80% HPDI: 0.02–0.14) and was 0.02 (80% HPDI: 1.10^-4^–0.04) during winter. The estimated contribution of juveniles to the effective reproduction number (*Re*) was approximately two orders of magnitude greater than in adults after birth pulse. Full posterior distributions of parameter estimates are provided in electronic supplementary material 2 (figures S4–S11; table S3), along with SEID/SER compartment dynamics (figure S12).

Overall mortality attributable to EBLV-2 infection was relatively low across the study period, averaging 5% in adults (80% HPDI: 0–13) and 4.4% in juveniles (80% HPDI: 0–12). However, during periods of elevated prevalence, the contribution of EBLV-2 to mortality increased substantially (figure 3c). Notably, during outbreak periods (2010 and 2014–2018), EBLV-2 infection was estimated to account for more than 50% of juvenile deaths, while in adults it became the leading cause of mortality (figure 3c; electronic supplementary material 2 figure S13). External FOI varied between 0 to more than 0.6 in 2014 and around 0,5 in 2015, 2016 indicating important contamination of the studied population from imported cases in 70% of the scenarios (figure S11 & Table S3, electronic supplementary material 2). The last estimate of the FOI is not identifiable, as its estimation in one year is conditioned by the prevalence in the subsequent year.

## Discussion

A robust, evidence-based understanding of how zoonotic pathogens transmit, persist, and impact host populations is essential for informing One Health strategies and conservation policies. In this study, we leverage longitudinal CMR data combined with sparse serological data to disentangle how key features of the host life cycle shape EBLV-2 transmission dynamics. Our analyses narrow the range of plausible epidemiological mechanisms that are consistent with the observed data. The inferred infection profile is characterised by relatively short-lasting immunity, short infectious periods, and a very low probability of lethal outcomes following exposure. These results support the view that EBLV2 is immunologically controlled by most individuals, such that epidemiological dynamics are driven by a small proportion of bats which suffer lethal infections. Transmission appears weak but highly structured, with consistently low *R_0_* values and strong support for age-dependent heterogeneity, highlighting the important role of juveniles in sustaining transmission. The duration of maternally derived immunity emerged as a key, but still poorly understood, factor which shaped whether the EBLV-2 maintenance relied on external reinfection or was maintained as enzootic infection. Short-lived maternal immunity, coupled with the birth pulse, synchronised the early recruitment of susceptible individuals during the rearing season, promoting high transmission peaks during mating season and cyclic epidemics, especially in the case of density-dependent transmission. In contrast, prolonged maternal immunity under frequency-dependent transmission delayed susceptibility until late in the mating season, increasing the likelihood of transmission-chain collapse and making external introductions necessary to sustain infection peaks. These findings emphasise the central role of early-life immunity in structuring transmission landscapes and indicate that relatively small shifts in immune protection can lead to qualitatively distinct epidemiological regimes. In this context, our results do not resolve the long-standing debate regarding density-versus frequency-dependent transmission in lyssaviruses [52], instead suggesting that maternal immunity duration may modulate the operation of these processes. Empirically determining the existence and duration of maternal immunity is therefore a main obstacle to identifying the precise mechanisms of transmission and persistence of EBLV-2. While observational studies of wild bats may in principle detect antibodies in young bats, which would suggest maternal immunity, such data do not exclude early life virus exposure. Antibody kinetics derived from longitudinal sampling of juveniles, from early life stages to several months of age, could theoretically help distinguish maternally derived immunity, which is expected to decline over time, from new infections, in which antibody titres typically increase and then decline more gradually. Studies in captive bats should be carried out to verify antibody transfer from mothers to pups and to experimentally assess the duration of protection following experimental lyssavirus challenge, but it is not feasible in a protected species like *M. myotis*.

Bat behaviour also emerged as a critical driver of viral dissemination. The mating season, during which multiple colonies and species aggregate, creates conditions conducive to transmission, particularly in the presence of newly recruited susceptible juveniles. Connectivity among colonies, which operate mainly during seasonal aggregations, in particular during the mating season, likely ensures viral spread across the network of colonies in scenarios where sporadic transmission explains the observed serological pattern. While earlier studies have often relied on simulations or strong assumptions, relatively few have fitted mechanistic models to multi-year serological data. Those that have, across different lyssaviruses [4,10,24,25], highlight the importance of bat movement and inter-colony or interspecific interactions in sustaining transmission, a conclusion further supported here.

The presence of EBLV-2 in different *M. myotis* colonies at large scale may follow a metapopulation structure, with source and sink populations whose roles vary in time and space. It is also possible that *M. myotis* itself functions as a sink with respect to other species such as *M. daubentonii*, which co-occur and share mating sites. In fact, the low infection rate in *M. myotis* may indicate frequent transmission or infection failures, consistent with a virus that is poorly adapted to this host species, either because early control of the virus may prevent it from reaching the brain by immune inactivation (interferon or pre-existing antibody) or the virus may be poorly efficient at binding to neural receptors or may replicate poorly in this host. This could explain the historically low detection of EBLV-2 in this species, but this should be revisited in future experimental or observational studies. We also observed seroreversion within a year, a pattern consistent with observational and experimental studies in the rabies virus [47,53]. This finding suggests that multi-year seropositivity, which we also observed, is more likely to reflect recurrent introductions than prolonged circulation or multi-year antibody persistence following a single outbreak. Indeed, the estimated immune protection in our models lasted approximately seven months, further supporting this interpretation.

Taken together, our results highlight the interplay between social behaviour, immunity, and spatial structure in shaping transmission. Seroprevalence patterns appear tightly linked to movement and contact among colonies and eventually species, peaking during the mating periods. These processes generate spatially mediated transmission dynamics characterised by localised and temporally variable waves of infection centred around mating sites. Consequently, connectivity networks and contact frequency emerge as key drivers of viral spread, intrinsically linking transmission to the mating system of *M. myotis*.

Bat lyssaviruses are not generally associated with large-scale mortality, yet some strains, including EBLV-1, have been linked to marked mortality events [54]. By integrating mortality estimates with mechanistic modelling, we show that infection-related mortality is low, affecting only a very limited number of individuals over the full study duration (figure S13, electronic supplementary material 2). Even when infection accounts for a substantial fraction of deaths, its relative contribution declines sharply during periods when other mortality drivers are operating in the meta-population (figure 3). The elevated mortality observed between 2012 and 2014, despite near-zero seroprevalence, was attributed to adverse weather conditions, particularly successive autumns, winters, and springs marked by cold temperatures and rainfall. During the peak of infection prevalence, the relative contribution of infection to mortality was higher in adults than in juveniles (figure 3); however, the absolute number of deaths remained low in both cases, with an average of fewer than 10 individuals in each class. Given that baseline mortality in juveniles was consistently high throughout the study, with approximately half of juveniles dying within their first year (Figure S4, Electronic Supplementary Material 2), the additional impact of EBLV-2 infection appears limited relative to other mortality sources. In this context, infection-associated mortality contributes variability rather than acting as a dominant force shaping population dynamics. Overall, EBLV-2 does not emerge as a primary driver of population-level mortality, but rather as one of several contributing factors.

Seropositivity was modelled as a binary variable, as it is generally the case in the field, however, quantitative titres could provide additional information (particularly the stage of the infection), but our lack of reference ranges prevents us from interpreting these values. Additionally, potential cross-reactivity with uncharacterised lyssaviruses cannot be excluded, given the diversity of viruses associated with *Myotis* bats [55].

This study identifies several avenues for future work to fully resolve the epidemiological dynamics of EBLV-2 and the maintenance mechanisms operating within and among bat populations and species. In particular, the duration of maternal immunity remains a key but difficult parameter to estimate, as discussed above. Annual and uneven sampling limits the ability to resolve intra-annual dynamics, and small sample sizes in some years likely contribute to discrepancies between observed and predicted seroprevalence. Increasing sampling intensity would improve parameter estimation and potentially allow colony-level modelling [4,48,56], although this must be balanced against disturbance, especially in this protected species. Finally, the absence of viral RNA detection despite substantial sampling effort is consistent with previous studies and reflects the rarity of detectable shedding [10]. This makes it difficult to detect and characterise the virus [57]. To establish antibody dynamics profiles, it would be necessary to carry out experimental infection studies on captive individuals monitored over the long term, which is not an option in the case of protected species [55,58].

From a public health perspective, the risk of human exposure to EBLV-2 from *M. myotis* in Brittany appears low, given the short infectious period, low prevalence of shedding, and sporadic dynamics. Human infections with bat lyssaviruses in Europe remain exceedingly rare, with only five confirmed cases since 1954, including two attributed to EBLV-2 [59]. Risk to the general public is minimal and largely limited to accidental contact, for which standard precautions and post-exposure prophylaxis are effective. However, attention is increasingly turning to domestic cats as potential bridge hosts. Their growing populations, frequent interactions with bats, and expanding overlap with bat habitats raise the risk of spillover, as documented in France and elsewhere [60,61]. The evidence of (albeit low) EBLV-2 circulation in *M. myotis* bats provided here implies a risk to cat owners and veterinarians and supports the need for awareness-raising and the systematic inclusion of lyssaviruses in the diagnostic work-up of cats presenting with fatal neurological signs. In conclusion, beyond introducing an integrated modelling framework, this study provides new insights into the mechanisms governing EBLV-2 transmission and persistence. Transmission is tightly constrained in space and time, largely occurring at mating sites during late summer and early autumn. While seasonal behaviours such as aggregation in maternity colonies limit inter-colony mixing, mating facilitates viral spread across connected colonies, generating transmission waves. Low *R_0_*, and limited infection success collectively preclude long-term persistence within isolated colonies but more probably external reintroduction through inter-colony or interspecific contacts. As a result, EBLV-2, like other lyssaviruses, is most likely maintained at the scale of a spatially structured metapopulation, despite low overall prevalence. More broadly, this work significantly advances the field by integrating immunological, demographic, and behavioural processes into a coherent framework for understanding pathogen persistence in wildlife systems.

### Ethics

Bat sampling was conducted in compliance with French ethical and sampling guidelines, under prefectural decrees issued by the Préfet du Morbihan (Brittany, France) of the July 9, 2010 for the period 2010-2012, of July 10, 2013 for the period 2013-2016 and of the May 5, 2017 for the period 2017-2020.

### Data and code accessibility

The data and the R codes used in this article are provided in the electronic supplementary material at https://zenodo.org/records/22300483.

### Authors’ contributions

FT: data collection, conceptualization, data curation, formal analysis, funding acquisition, investigation, methodology, validation, visualization, writing original draft. MW: conceptualization, data curation, methodology, validation, writing-review and editing.

EP: conceptualization, data curation, methodology, validation, supervision, writing-review and editing.

OF, YL, AL, CLF, SJP: data collection, conceptualization, investigation, methodology, writing-review and editing.

MEG: formal analysis, writing-review and editing.

ET: data collection, conceptualization, funding acquisition, validation, visualization, supervision, writing-review and editing.

JM, MV: formal analysis, validation, visualization, writing-review and editing DGS: conceptualization, supervision, writing-review and editing

All authors gave final approval for publication and agreed to be held accountable for the work performed therein.

### Funding

F.T. has received funding for this research from the European Union’s Horizon 2020 research and innovation programme under the Marie Skłodowska-Curie grant agreement No 101034345. This research was supported by a Science Foundation Ireland (SFI) Future Frontiers Award 19/FFP/6790, a European Research Council Starting (Grant 2012-StG311000) and Synergy grant (no. 101118919) awarded to ECT. D.S. was supported by a Wellcome Trust Senior Research Fellowship (217221/Z/19/Z) and the UK Biotechnology and Biological Sciences Research Council BBSRC (BB/V003798/1). Additional support for this work was provided by UK Medical Research Council (MRC) through core support to the MRC-University of Glasgow Centre for Virus Research (Virus Cross Species Transmission Programme: MC\UU\00034/3).

## Acknowledgements

We want to acknowledge all the dedicated volunteers of the regional conservation NGO Bretagne Vivante, whose unwavering commitment to monitoring and capturing bats made this long-term study possible, as the students and volunteers of the UCD bat lab that aided in the sample collections throughout the years. We are also deeply grateful to Dan Haydon for his insightful and inspiring discussions on the modelling approach during the early stages of this manuscript.

---

**Table 1.**
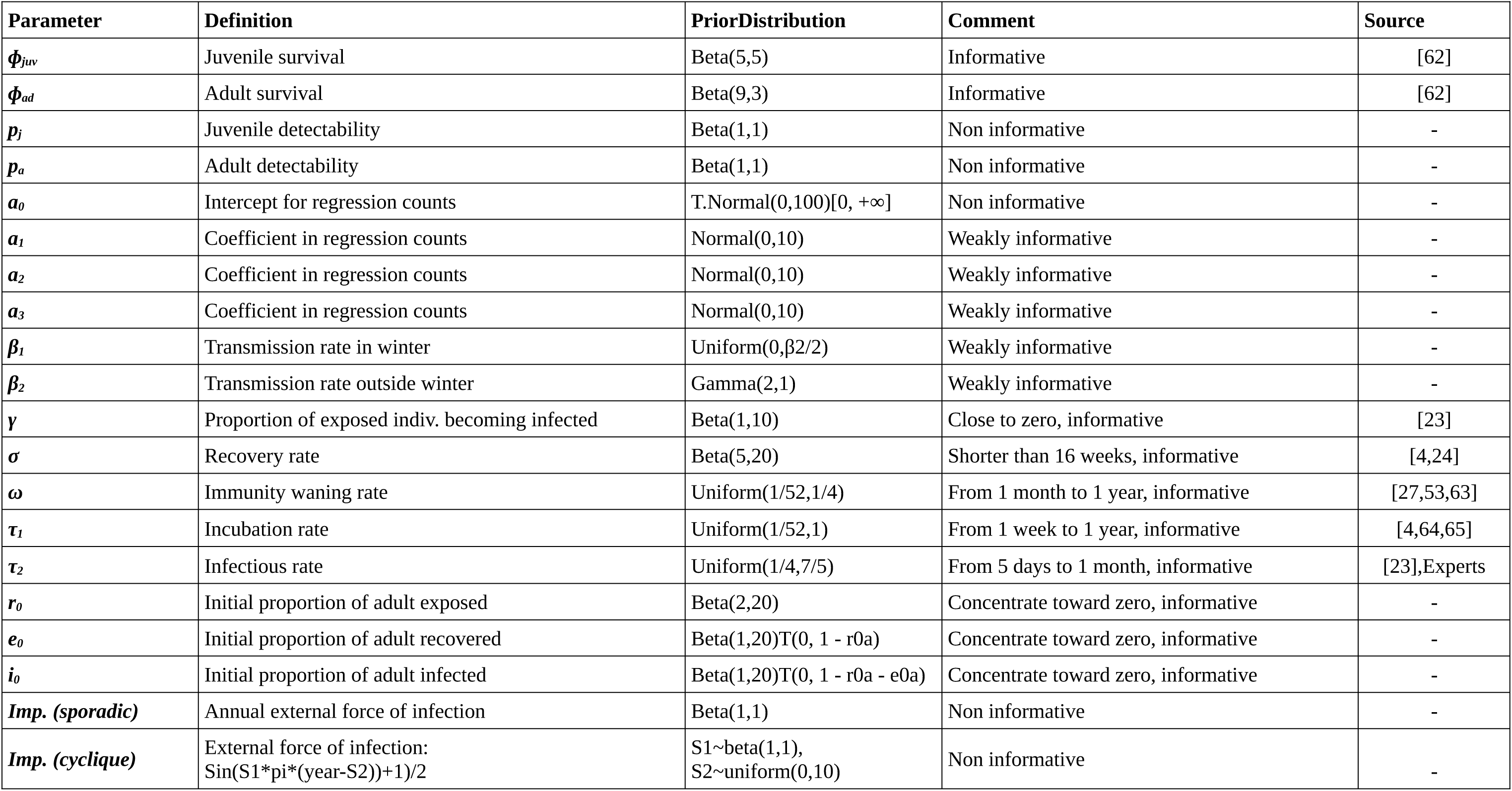
Priors definition and specification.

| Parameter | Definition | PriorDistribution | Comment | Source |
| --- | --- | --- | --- | --- |
| $\phi_{juv}$ | Juvenile survival | Beta(5,5) | Informative | [62] |
| $\phi_{ad}$ | Adult survival | Beta(9,3) | Informative | [62] |
| $p_j$ | Juvenile detectability | Beta(1,1) | Non informative | - |
| $p_a$ | Adult detectability | Beta(1,1) | Non informative | - |
| $a_0$ | Intercept for regression counts | T.Normal(0,100)[0, $+\infty$] | Non informative | - |
| $a_1$ | Coefficient in regression counts | Normal(0,10) | Weakly informative | - |
| $a_2$ | Coefficient in regression counts | Normal(0,10) | Weakly informative | - |
| $a_3$ | Coefficient in regression counts | Normal(0,10) | Weakly informative | - |
| $\beta_1$ | Transmission rate in winter | Uniform(0,$\beta_2/2$) | Weakly informative | - |
| $\beta_2$ | Transmission rate outside winter | Gamma(2,1) | Weakly informative | - |
| $\gamma$ | Proportion of exposed indiv. becoming infected | Beta(1,10) | Close to zero, informative | [23] |
| $\sigma$ | Recovery rate | Beta(5,20) | Shorter than 16 weeks, informative | [4,24] |
| $\omega$ | Immunity waning rate | Uniform(1/52,1/4) | From 1 month to 1 year, informative | [27,53,63] |
| $\tau_1$ | Incubation rate | Uniform(1/52,1) | From 1 week to 1 year, informative | [4,64,65] |
| $\tau_2$ | Infectious rate | Uniform(1/4,7/5) | From 5 days to 1 month, informative | [23],Experts |
| $r_0$ | Initial proportion of adult exposed | Beta(2,20) | Concentrate toward zero, informative | - |
| $e_0$ | Initial proportion of adult recovered | Beta(1,20)T(0, 1 - r0a) | Concentrate toward zero, informative | - |
| $i_0$ | Initial proportion of adult infected | Beta(1,20)T(0, 1 - r0a - e0a) | Concentrate toward zero, informative | - |
| *Imp. (sporadic)* | Annual external force of infection | Beta(1,1) | Non informative | - |
| *Imp. (cyclique)* | External force of infection:<br>$\text{Sin}(S1*\pi*(\text{year}-S2))+1)/2$ | $S1\sim\text{beta}(1,1)$,<br>$S2\sim\text{uniform}(0,10)$ | Non informative | - |

